# CSF1R regulates monocyte subset differentiation and intracellular metabolism

**DOI:** 10.1101/2025.07.17.665275

**Authors:** Alexandre Gallerand, Johanna Merlin, Zakariya Caillot, Chloé Delaby, Elisa Bord, Jichang Han, Bastien Dolfi, Alexia Castiglione, Sacha Grenet, Maxime Franceschini, Gisèle Jarretou, Fairouz N. Zair, Evy Boré, Florian Tuffin, David Dombrowicz, Rodolphe R. Guinamard, Gwendalyn J. Randolph, Adeline Bertola, Patrick Auberger, Arnaud Jacquel, David A. Hume, Jesse W. Williams, Marc Bajénoff, Jaap G. Neels, Stoyan Ivanov

**Author notes:** Contributed equally (Alexandre Gallerand, Johanna Merlin). Co-last authors (Jaap G. Neels, Stoyan Ivanov).

## Abstract

Monocytes are key circulating effectors of vascular homeostasis, innate immunity and inflammation. Following their generation in mouse bone marrow, classical (Ly6C^high^) monocytes are mobilized into the blood circulation where they mature into non-classical (Ly6C^low^) patrolling monocytes or are recruited into peripheral tissues where they differentiate into tissue resident or inflammatory macrophages. Monocytes and macrophages express CSF1R (CD115), the receptor for lineage-specific growth factors CSF1 and IL34. Here, we report that acute CSF1R blockade or genetic deletion negatively interferes with monocyte intracellular metabolism and reduces blood Ly6C^low^ monocytes in part by blunting differentiation of Ly6C^high^ monocytes. Based upon lineage-specific deletion of GFPT1 (Glutamine-Fructose-6-Phosphate Transaminase 1), the hexosamine biosynthetic pathway (HBP) is identified as a novel regulator of CSF1R expression and monocyte subset diversity. Our findings provide new insights into the link between CSF1R signaling, metabolic regulation, and monocyte survival and differentiation.

## Introduction

Monocytes are innate immune cells generated in bone marrow by a tightly and highly regulated process named myelopoiesis ^1^. Multiple consecutive steps lead to monocyte generation from hematopoietic stem cells (HSCs) through several intermediate committed myeloid progenitors ^1^. Among those, granulocyte/macrophage progenitor (GMPs) have been shown to give rise to the majority of blood circulating monocytes ^2^. In mouse blood, two major monocyte subsets have been identified. Ly6C^high^ (classical monocytes) that express the chemokine receptor CCR2 infiltrate into peripheral tissues and contribute to tissue macrophage pool size at steady-state and upon inflammation ^3-5^. Less is known about the generation and functions of Ly6C^low^ monocytes, previously shown to patrol blood endothelial cells and providing growth factors ^6,7^. While most blood monocytes originate from GMPs, recent evidence indicates that monocyte/dendritic progenitors (MDPs) also generate a defined subset of blood monocytes ^2,8^.

Hematopoietic stem cell commitment to the monocyte-macrophage lineage is associated with inducible expression of *Csf1r,* encoding the receptor for macrophage colony-stimulating factor (CSF1) and a second ligand, IL34 ^9^. CSF1R protein is a lineage-restricted marker, expressed by all committed progenitors, blood monocytes and resident tissue macrophages ^10-12^. CSF1R (CD115) is a class III receptor tyrosine kinase (RTK) structurally related to FLT3 (CD135) and KIT (CD117), which are also expressed by myeloid progenitors. CSF1 promotes the proliferation and differentiation of mouse bone marrow cells or human monocytes *in vitro* to produce bone marrow-derived macrophages, or monocyte-derived macrophages, widely-used models for the study of macrophage biology ^13^. *In vivo*, administration of recombinant CSF1, or a CSF1-Fc fusion protein to mice, causes a profound and selective monocytosis and proliferative expansion of tissue resident macrophage populations ^14-16^. Mutations in the *Csf1* and *Csf1r* genes in mice and rats lead to severe osteopetrosis (osteoclast deficiency) and the loss of many tissue resident macrophage populations ^17,18^. Notwithstanding these observations, multiple lines of evidence indicate that CSF1R signaling is not absolutely required for monocytopoiesis. Instead, treatment of mice with blocking anti-CSF1R or anti-CSF1 antibodies leads to selective loss of Ly6C^low^ non-classical monocytes ^19-21^. Similarly, inducible genetic deletion of *Csf1r* or endothelial *Csf1* affected Ly6C^low^ but not Ly6C^high^ monocytes ^22,23^. A *Csf1r* enhancer mutation that abolishes CSF1R expression and CSF1 responsiveness in bone marrow progenitors had no effect on blood monocyte numbers and subset diversity ^24^. Our recent work demonstrated that glucose metabolism sustains monocyte CSF1R expression, and inhibiting glucose uptake was associated with decreased bone marrow, blood and spleen monocyte numbers ^25^. In this study we explore the mechanisms that connect CSF1R signaling, glucose metabolism and monocyte differentiation.

## Results

### Epitope masking confounds identification of blood monocytes via CSF1R expression

Previous studies documented that the administration of anti-CSF1R (CD115) antibodies to mice robustly reduced blood monocyte counts, a phenotype sometimes observed on both Ly6C^high^ and Ly6C^low^ monocytes ^26,27^ or solely on Ly6C^low^ monocytes^20-22,28^. We also observed that downregulation of CSF1R precedes loss of blood monocytes ^25^. To explore causal links between CSF1R expression and monocyte survival, mice received two intraperitoneal (i.p.) injections of anti-CSF1R (AFS98) or isotype control (rat IgG2a) 48 hours apart and were analyzed 24 hours later (**Figure 1A**). This regimen was effective in depletion of white adipose tissue (WAT) macrophages. CSF1R-dependent CD206^high^ WAT macrophages, identified as CD45^+^ CD206^+^ MerTK^+^ cells, were robustly depleted in AFS98-injected mice as reported previously ^29^ (**Figure 1A**). Blood monocytes were initially analyzed using a gating strategy relying on the combination of CD11b and CSF1R staining (**Figure 1B**). In comparison to isotype control-treated mice, an almost complete disappearance of CD11b^+^ CSF1R^+^ cells was observed in AFS98-injected animals (**Figure 1B**). We repeated this experiment in a time-course manner and observed that CD11b^+^ CSF1R^+^ cells were undetectable 1 hour post AFS98 injection and started re-appearing 72 hours after injection (**Figure 1C**). Since CSF1R blockade leads to serum accumulation of CSF1 ^16,21,28^, blood was drawn from C57BL/6 mice, kept on ice to prevent receptor internalization, then incubated with or without CSF1 for 45 minutes, washed and stained for CSF1R using PE-conjugated AFS98 antibody (**Figures 1D and S1A**). CD115 staining declined in a dose-dependent manner after incubation with recombinant CSF1 *ex vivo* (**Figure 1D**), confirming that CSF1 directly prevented AFS98 binding.

**Figure 1.**
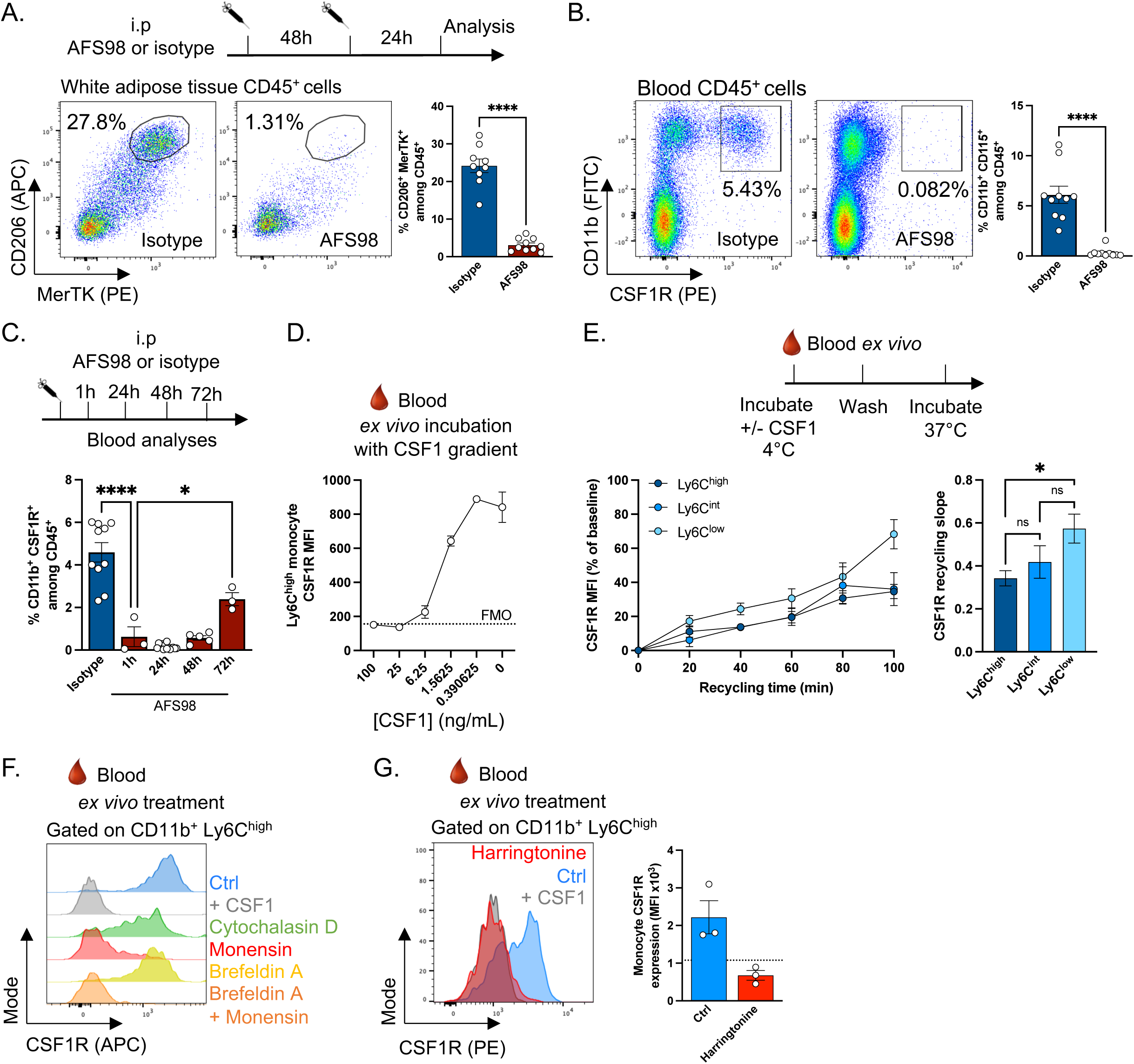
CSF1R epitope masking by AFS98 and CSF1. (**A**) Experimental scheme used to test impact of AFS98 administration on blood monocytes (top). Representative flow cytometry plots and proportions of CD206^+^ MerTK^+^ adipose tissue macrophages following AFS98 treatment (bottom). (**B**) Representative flow cytometry plots (left) and proportions (right) of CD11b^+^ CSF1R^+^ cells in blood following AFS98 treatment. (**C**) Expression of CSF1R detected through AFS98-PE conjugated antibody staining 1, 24, 48 or 72 hours after injections of 500μg AFS98. (**D**) Detection of CSF1R using AFS98-PE conjugated antibody after incubation with increasing concentrations of recombinant CSF1. (**E**) Experimental scheme used to measure CSF1R renewal after CSF1 binding (top) and measure of CSF1R expression on blood monocyte subsets using AFS98-APC conjugated antibody in a time-course manner after incubation at 37°C to allow receptor recycling (bottom). (**F-G**) Measure of CSF1R expression on blood monocytes using PE-conjugated AFS98 after a 45 minutes incubation at 37°C with Cytochalasin D, Monensin, Brefeldin A (F) or Harringtonine (G).

CSF1R is rapidly internalized following ligand binding, ubiquitinated and degraded ^17,30^. The response to CSF1 analyzed in bone marrow-derived macrophages suggests that macrophages require continuous resynthesis of the receptor and its reappearance on the cell surface. The rate of receptor synthesis can be monitored by flow cytometry in cells in which CSF1-repleted cells are washed free of ligand ^31-33^. To determine CSF1R recycling rate in monocytes, we incubated blood with recombinant CSF1 at 4°C to mask surface epitopes and then washed the cells thoroughly to remove any potentially unbound CSF1 protein. Cells were then kept on ice (baseline) or incubated at 37°C (without CSF1) for up to 100 minutes to allow the turnover of the receptor (**Figure 1E**). Surface CSF1R expression gradually reappeared over time in Ly6C^high^ Treml4^-^ (Ly6C^high^), Ly6C^+^ Treml4^+^ (Ly6C^int^) and Ly6C^low^ Treml4^+^ (Ly6C^low^) monocytes (**Figure 1E and Supplementary Figure 1B**). Importantly, CSF1R recycling rate was higher in Ly6C^low^ compared to Ly6C^high^ monocytes (**Figure 1E**).

To explore mechanisms regulating surface CSF1R expression, we first inhibited actin polymerization using cytochalasin D to broadly impair intracellular protein trafficking. This resulted in a partial downregulation of CSF1R expression from the cell surface after only 45 minutes, indicating that CSF1R is continuously exported to the cell surface to maintain its expression (**Figure 1F**). Inhibiting Golgi-dependent trafficking with monensin led to a strong reduction of surface CSF1R signal, while impairing endoplasmic reticulum (ER) to Golgi trafficking with brefeldin A only had a partial effect (**Figure 1F**). A combined treatment with monensin and brefeldin A completely abrogated surface CSF1R expression on monocytes (**Figure 1F**). Inhibition of protein translation with harringtonine also rapidly reduced CSF1R surface expression on blood monocytes (**Figure 1G**). Together, this set of results demonstrated that blood monocyte surface CSF1R expression requires continuous protein translation and actin-dependent addressing to the cell membrane following maturation in the Golgi apparatus, rather than receptor endocytosis and recycling.

### CSF1R sustains Treml4^+^ monocyte survival and commitment into Ly6C^low^ monocytes

To avoid the use of CD115 as a marker, we devised an alternative gating strategy. Monocytes were defined as CD45^+^ CD11b^+^ NK1.1^-^ Ly6G^-^ SSC^low^ cells then further divided into Ly6C^high^ (R1) and Ly6C^low^ (R2) subsets (**Figure 2A**). Using CX3CR1^gfp^ mice ^34^ we observed a robust expression of CX3CR1 on both R1 and R2 cells, with the latter showing higher and homogenous expression of the reporter in comparison to R1 cells (**Figure 2B**). In contrast, using CCR2^cre/ERT2^ x R26TdT^LSL^ mice ^35^, we detected higher TdTomato expression in the R1 population compared to R2 (**Figure 2B**). Both R1 and R2 cells were CD115-positive in untreated control mice (**Figure 2B**). **Figure 2C** compares the relative frequency of R1 and R2 cells in isotype control and AFS98-treated mice. As reported by others ^20^, AFS98 administration had no effect on blood Ly6C^high^ monocytes but selectively reduced Ly6C^low^ monocytes (**Figure 2C**). These data demonstrated that Ly6C^low^ monocytes, but not Ly6C^high^ monocytes, are particularly sensitive to CSF1R blockade.

**Figure 2.**
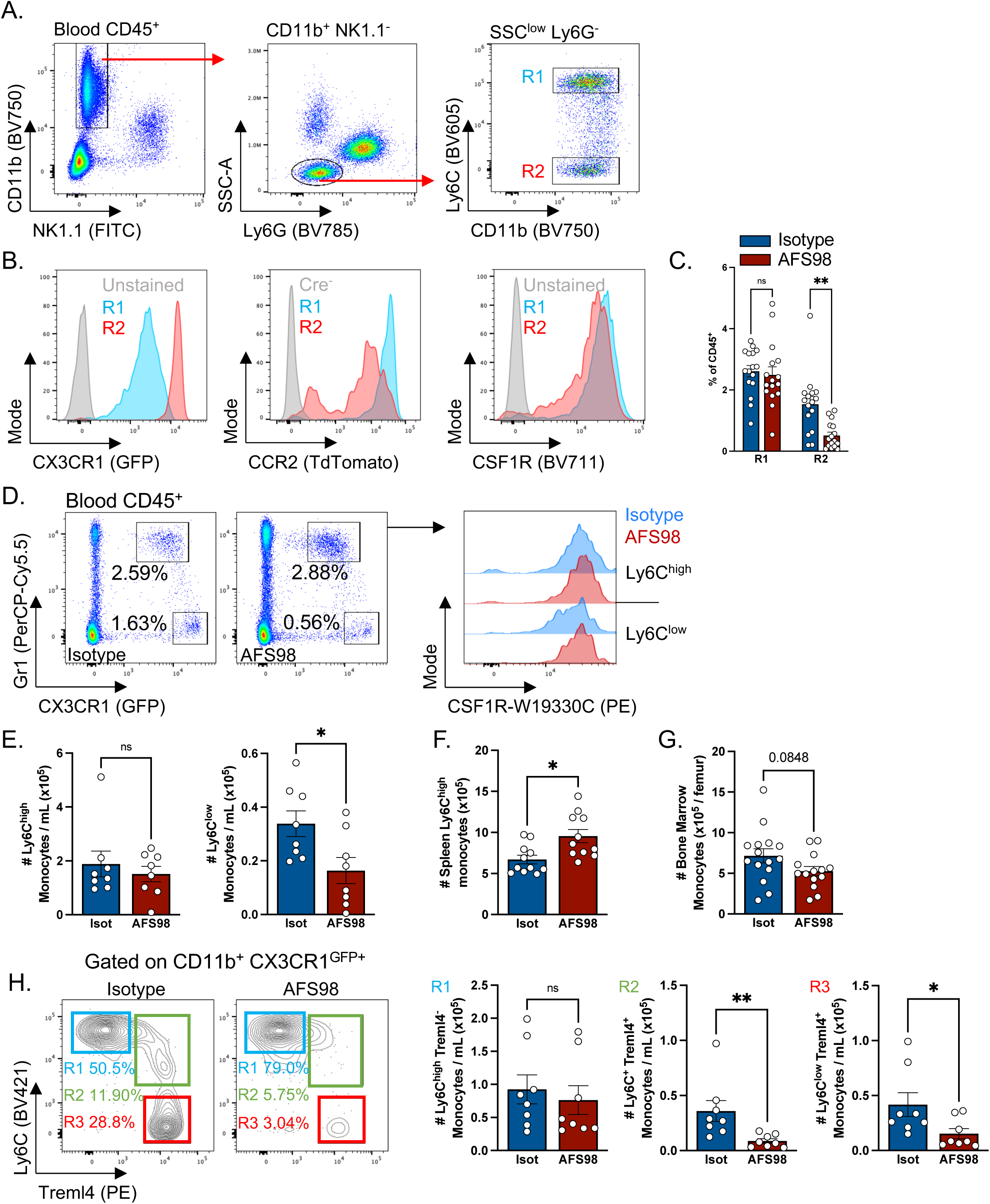
CSF1R blockade impairs Treml4+ monocytes. (**A**) Gating strategy used to identify Ly6C^high^ (R1) and Ly6C^low^ (R2) cells without including CSF1R. (**B**) Histograms representing expression of CX3CR1^gfp^, CCR2^TdTomato^ and CSF1R by R1 and R2 cells. (**C**) Proportions of R1 and R2 cells among total immune cells following AFS98 treatment. (**D**) Representative flow cytometry plots illustrating presence of CSF1R^+^ Gr1^high^ CX3CR1^+^ and CSF1R^+^ Gr1^low^ CX3CR1^high^ blood cells following treatment with AFS98 or isotype control. (**E-G**) Quantification of blood (E), spleen (F) and bone marrow (G) monocytes identified as CX3CR1^GFP+^ CSF1R^+^ (probed with W19330C-PE conjugated antibody) in CX3CR1^GFP/+^ mice that received isotype or AFS98 treatment as in Figure 1A. (**H**) Gating strategy applied to analyze blood monocyte subsets in AFS98 and isotype control treated mice.

The most obvious explanation for the loss of detectable CD115 in mice treated with AFS98 is that monocytes are saturated with unlabeled antibody. To address this question, we tested alternative anti-CD115 antibodies. CSF1R signal was reduced after AFS98 injection when CSF1R was probed using a sheep polyclonal antibody (AF3818) or monoclonal clones REA827 and W19330E (**Supplementary Figure 1C**). However, CSF1R signal was retained in AFS98-treated mice when detected using monoclonal antibody W19330C (**Supplementary Figure 1C**). The specificity was confirmed in that incubation with CSF1 inhibited CSF1R detection with all clones (**Supplementary Figure 2A**).

Based on these new findings, we repeated *in vivo* CSF1R blockade (as described in **Figure 1A)** in CX3CR1^GFP/+^ mice, and quantified blood monocyte numbers, identified through co-expression of GFP and CSF1R probed using the W19330C clone (**Figure 2D**). In accordance with results obtained using our CSF1R-independent gating strategy (**Figure 2A-C**), Ly6C^high^ monocyte numbers were unaffected by CSF1R blockade, while Ly6C^low^ monocyte numbers were diminished by half (**Figure 2E**). We used Uniform Manifold Approximation and Projection (UMAP) dimensional reduction of spectral flow cytometry data to obtain a clearer picture of monocyte phenotype following AFS98 Ab administration (**Supplementary Figure 2B**). General UMAP distribution of CD11b^+^ CX3CR1^gfp+^ cells from isotype-treated and AFS98-treated mice was comparable (**Supplementary Figure 2B**). Ly6C^high^ and Ly6C^low^ cells were both detectable with the latter expressing canonical patrolling monocyte markers CD43 and Treml4, as well as CD11c (**Supplementary Figure 2C**) ^10,11,36^ and being depleted in AFS98-injected mice (**Supplementary Figure 2C**). We also noticed that monocytes with maximal expression of Ly6C appeared in AFS98-treated mice (**Supplementary Figure 2C**), potentially indicating that monocyte egress from bone marrow was accelerated or differentiation to Ly6C^low^ monocytes was halted by CSF1R blockade. We detected an accumulation of Ly6C^high^ monocytes in the spleen of AFS98-treated mice (**Figure 2F**). Since CSF1R is expressed on bone marrow and spleen monocyte progenitors ^37^, we aimed to determine whether these phenotypes could be due to altered monocyte generation or mobilization. Bone marrow monocyte numbers tended to be diminished (**Figure 2G**), while neutrophils and myeloid progenitors were unaffected (**Supplementary Figures 2D-E**).

Following prolonged treatment with an alternative blocking anti-CSF1R antibody, the loss of Ly6C^low^ monocytes was balanced by an increase in Ly6C^high^ monocytes implying inhibition of conversion as a mechanism (Macdonald et al., 2010). We tested this suggestion in more detail using the acute treatment regime with AFS98. Ly6C^high^ Treml4^-^ monocytes were not affected by AFS98 administration (**Figure 2H**). However, the numbers of Ly6C^+^ Treml4^+^ monocytes were reduced, ultimately culminating in decreased counts of Ly6C^low^ monocytes (**Figure 2H**). Together these data provide further evidence that monocytopoiesis is CSF1R-independent ^19^ whereas monocyte differentiation into the Ly6C^low^ subset is affected by CSF1R blockade. Thus, the decreased frequency and numbers of Ly6C^low^ monocytes likely results from two complementary processes: 1) diminished commitment of Ly6C^+^ Treml4^+^ monocytes; 2) reduced circulating half-life of Ly6C^low^ Treml4^+^ cells described previously in mice lacking CSF1 expression in endothelial cells ^23^.

### CSF1R differentially regulates Ly6C^low^ monocytes depending on their origin from cMoPs and MDPs

The vast majority of blood monocyte are generated from GMP ^2^ but a subpopulation derived from MDP can be distinguished by surface markers ^38^. We identified and validated Clec9a^iCre^ mice as a novel tool to target and analyze MDP-derived monocytes ^29^. We generated Clec9a^iCre^ x R26TdTomato^LSL^ mice and compared the phenotype of TdTomato^-^ (GMP-derived) and TdTomato^+^ (MDP-derived) monocytes. TdTomato^+^ cells were distinguished from TdTomato^-^ cells by increased CSF1R expression (**Figure 3A**). We administered AFS98 Ab to Clec9a^icre^ x R26TdTomato^LSL^ mice using the protocol described in **Figure 1A**. CSF1R blockade had no significant impact on bone marrow monocyte subsets (**Figure 3B**). However, the percentage of TdTomato^+^ Ly6C^low^ monocytes in blood and spleen was increased in AFS98-treated mice whereas TdTomato^+^ Ly6C^high^ monocytes were unaffected (**Figure 3C and 3D**). Finally, the relative resistance of MDP-derived monocytes to AFS98 treatment was not a direct consequence of a differential CSF1R turnover (**Figure 3E**). CSF1R reappearance at the cell surface was comparable between GMP- and MDP-derived monocytes (**Figure 3E**). Therefore, MDP-derived monocytes, while displaying higher CSF1R expression, appeared less dependent on CSF1R signaling.

**Figure 3.**
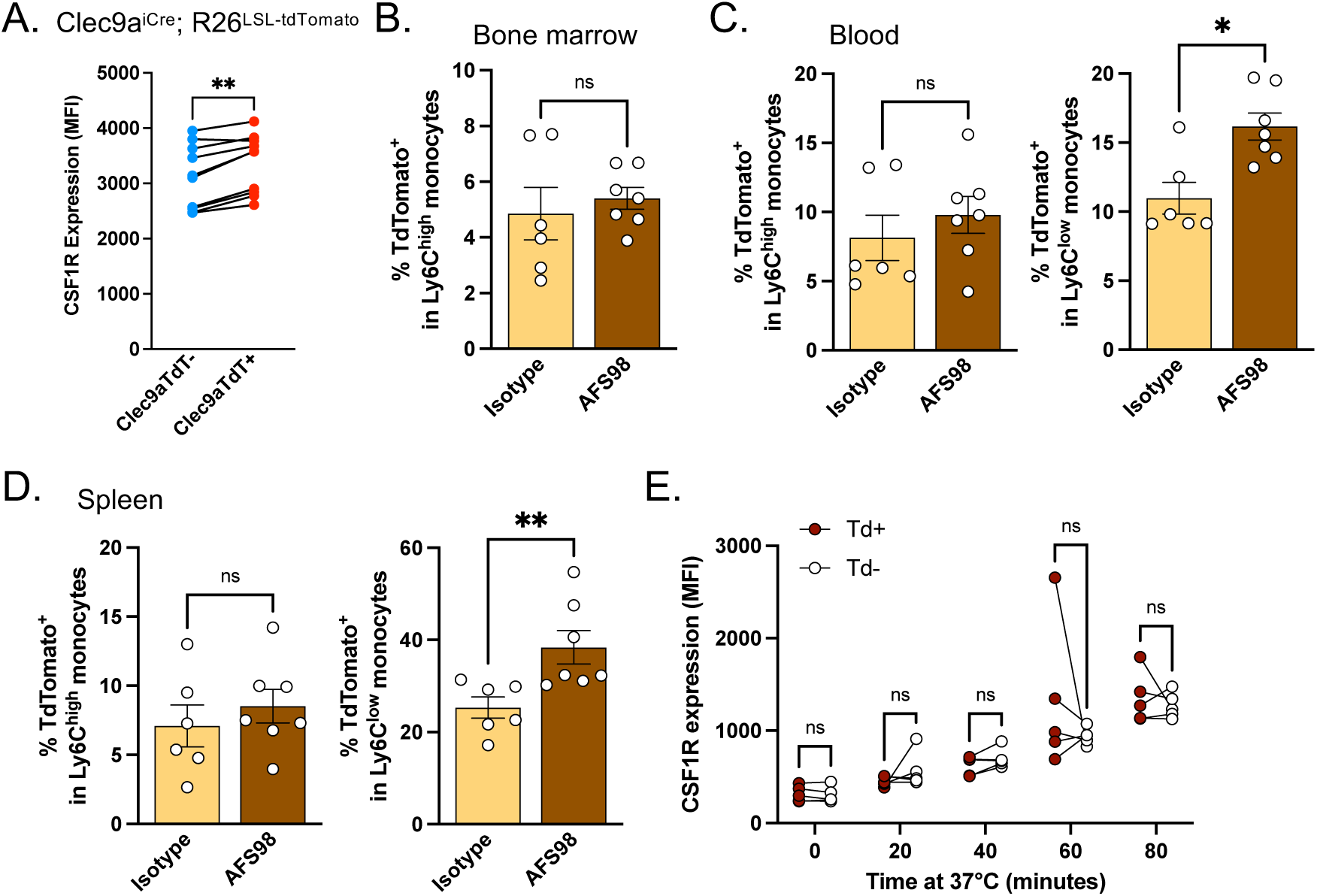
AFS98 treatment differentially affects cMoP- and MDP-derived monocytes. (**A**) Quantification of surface CSF1R expression in tdTomato^+^ and tdTomato^-^ blood monocytes from Clec9a^iCre^;R26^tdTomato^ mice. (**B**) Proportions of tdTomato^+^ cells among bone marrow monocytes. (**C and D**) Proportions of tdTomato^+^ cells among blood and spleen Ly6C^high^ and Ly6C^low^ monocytes from Clec9a^iCre^;R26^tdTomato^ mice treated with isotype control or AFS98 as in Figure 1A. (**E**) Analysis of CSF1R turnover in blood monocytes from Clec9a^iCre^;R26^tdTomato^ mice.

### CSF1R sustains Ly6C^low^ monocytes in Csf1r^ΔFIRE^ mice

The role of CSF1R in blood monocyte differentiation was explored further using genetic models. A conditional deletion model (CX3CR1^creERT2^ x R26^TdTomato^ x Csf1r^flox/flox^) is shown in **Figure 4A and Supplementary Figure 3A**. Following tamoxifen treatment, the proportions of TdTomato-labeled Ly6C^high^ monocytes tended to increase in blood, while TdTomato^+^ Ly6C^low^ monocyte proportions decreased (**Figure 4B**). We conducted similar analyses in Csf1r^ΔFIRE^ (FIRE/FIRE) mice, which were shown to lack CSF1R expression in bone marrow progenitors and are unresponsive to CSF1 due to deletion of an enhancer locus ^24^. Ly6C^low^ Treml4^+^ monocytes were enriched among total leukocytes in the blood of FIRE mice, while Ly6C^high^ monocytes tended to decrease (**Figure 4C**). A similar phenotype was observed in spleen from FIRE mice (**Figure 4D**), while their bone marrow monocytes were reduced (**Figure 4E**). These observations revealed that CSF1R affected monocyte homeostasis in FIRE mice. In the original description of these mice ^24^ CSF1R expression was detectable on Ly6C^low^ monocytes, while no signal was detected on Ly6C^high^ monocytes. This finding is confirmed in **Figure 4F**. To explore whether the remaining CSF1R expression on Ly6C^low^ monocytes could be functionally important, AFS98 Ab was injected in Csf1r^ΔFIRE/ΔFIRE^ and Csf1r^ΔFIRE/+^ littermate controls. The mutation is not dosage compensated so the heterozygotes have 50% reduced CSF1R ^24^. In both groups AFS98 Ab treatment completely depleted blood Ly6C^low^ monocytes (**Figure 4G**).

**Figure 4.**
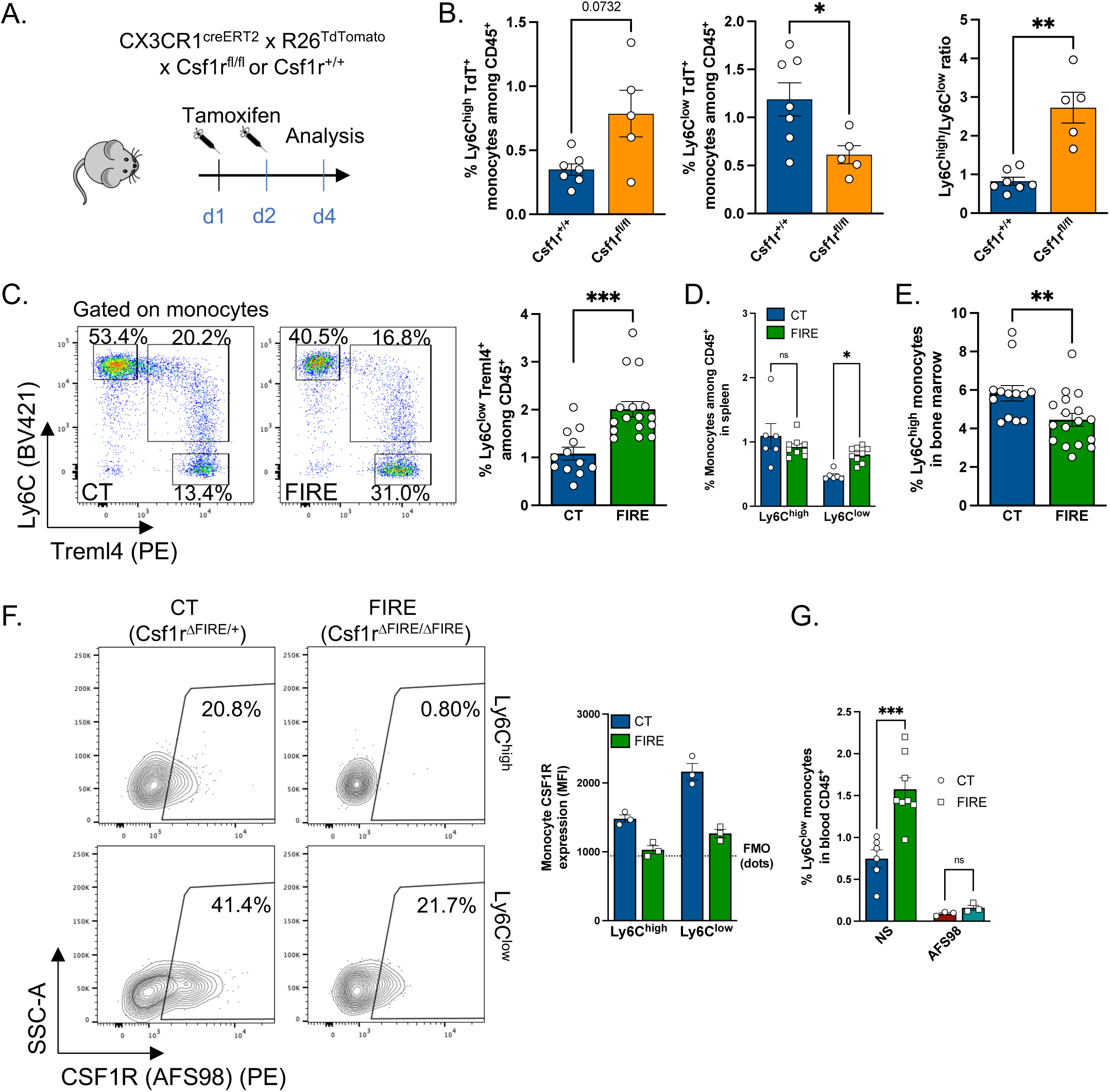
Ly6C^low^ monocytes rely on CSF1R signaling in FIRE mice. (**A**) Experimental scheme used to induce *Csf1r* deletion and tdTomato expression in CX3CR1^creERT2^ mice. (**B**) Proportions of Ly6C^high^ and Ly6C^low^ monocytes and ratio between these two subsets in CX3CR1^creERT2^ x Csf1r^fl/fl^ x R26^tdTomato^ mice and CX3CR1^creERT2^ x Csf1r^+/+^ x R26^TdTomato^ controls. (**C-E**) Proportions of monocytes in blood (C), spleen (D) and bone marrow (E) from Csf1r^ΔFIRE/ΔFIRE^ (FIRE) mice or their littermate Csf1r^ΔFIRE/+^ controls (CT). (**F**) Representative flow cytometry plots (left) and quantification (right) or CSF1R expression by blood Ly6C^high^ and Ly6C^low^ monocytes from FIRE and littermate CT mice. (**G**) Proportions of blood Ly6C^low^ monocytes in FIRE and littermate CT mice at steady state (NS) or following treatment with AFS98.

### The hexosamine biosynthetic pathway controls CSF1R expression and Ly6C^low^ monocyte homeostasis

In comparison to closely related myeloid cells such as macrophages and dendritic cells, data on the metabolic configuration of blood monocyte subsets in mice and humans are scarce ^39^. In monocyte-derived macrophages, CSF1 binding to CSF1R supports metabolic reprogramming towards glycolysis ^40,41^. Inhibiting glucose metabolism leads to a rapid and consistent downregulation of CSF1R on monocytes^25^. Therefore, we investigated intracellular metabolism pathways in blood monocytes after AFS98 Ab treatment. CSF1R blockade tended to inhibit AKT and mTOR phosphorylation in blood Ly6C^low^ but not Ly6C^high^ monocytes, reflecting an altered metabolic state (**Figure 5A and Supplementary Figure 3B**). We next applied the recently described SCENITH approach to define monocyte metabolic configuration ^42^. Although presenting several limitations, this approach monitors intracellular metabolism at a single-cell level resolution. Ly6C^high^ monocytes incorporated more glucose and puromycin than Ly6C^low^ cells (**Figure 5B and Supplementary Figure 3C**). AFS98 Ab treatment reduced glucose uptake in Ly6C^high^ monocytes (**Figure 5B**). Monocytes are highly dependent on glucose intracellular metabolism ^25^, independently of isotype or AFS98 treatment (**Figure 5C**). However, CSF1R blockade sensitized monocytes to oligomycin treatment, resulting in higher mitochondrial dependence and lower glycolytic capacity compared to control cells (**Figure 5C**). CSF1R signaling thus appears to directly regulate monocyte intracellular metabolism and dependency on glucose metabolism or mitochondrial oxidative phosphorylation. Consistent with evidence that CSF1R signaling is active in monocytes in FIRE mice, despite low receptor expression, monocyte metabolism was not significantly affected by the homozygous mutation (**Supplementary Figure 3D**).

**Figure 5.**
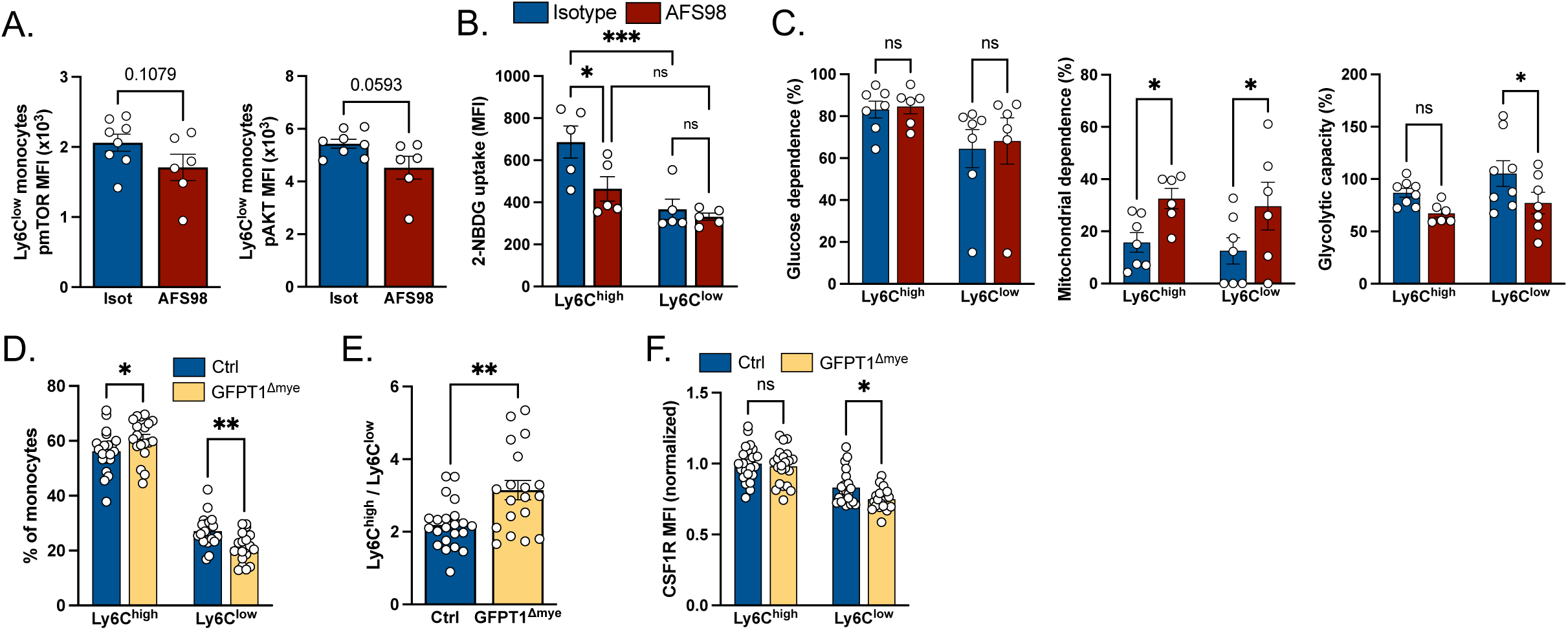
Regulation of monocytes by HBP. (**A**) Quantification of mTOR and AKT phosphorylation in blood Ly6C^low^ monocytes following CSF1R blockade or isotype control administration. (**B**) 2-NBDG incorporation in blood Ly6C^high^ and Ly6C^low^ monocytes following CSF1R blockade or isotype control administration. (**C**) Metabolic profile of blood monocytes following treatment with AFS98 or isotype control measured using SCENITH. (**D**) Frequency and **(E)** ratio of blood Ly6C^high^ and Ly6C^low^ monocyte populations in control and GFPT1^ΔMye^ mice. **(F)** CSF1R expression on Ly6C^high^ and Ly6C^low^ blood monocytes.

We previously demonstrated that glucose metabolism plays a central role in controlling myelopoiesis and monocyte numbers ^25^. We demonstrated that preventing glucose entry, by administrating a Glut1-inhibitor, dramatically decreased monocyte CSF1R expression and their numbers in peripheral blood ^25^. Once internalized, glucose is integrated and metabolized by 3 pathways: glycolysis, the pentose phosphate pathway (PPP) and the hexosamine biosynthetic pathway (HBP). Our previous data using genetic models suggested that glycolysis and PPP are not essential for CSF1R regulation ^25^. However, such a tool was not available to analyze the contribution of HBP and studies regarding this pathway heavily relied on pharmacological inhibitors, the most applied being tunicamycin.

GFPT1 (Glutamine-Fructose-6-Phosphate Transaminase 1) is the first and rate-limiting enzyme in HBP. Recently, GFPT1^fl/fl^ mice were generated and the ablation of this enzyme affected T cell thymic generation ^43^. To investigate the role of HBP in monocyte biology, with a particular focus on CSF1R expression, we generated Lyz2^cre^ x GFPT1^fl/fl^ mice (named GFPT1^ΔMye^) (**Supplementary Figure 4A**). Recombination efficiency in monocytes was confirmed by PCR on genomic DNA (**Supplementary Figure 4B**). Western blot analysis demonstrated the complete loss of GFPT1 protein in GFPT1^ΔMye^ bone marrow-derived macrophages in comparison to control cells (**Supplementary Figure 4C**). The frequency of Ly6C^high^ blood monocytes increased while that of Ly6C^low^ decreased in GFPT1^ΔMye^ mice in comparison to littermate control animals (**Figure 5D**), so that the ratio of Ly6C^high^/Ly6C^low^ blood monocytes was increased around 1.5-fold (**Figure 5E**). CSF1R expression was marginally decreased in GFPT1-deficient Ly6C^low^, but not Ly6C^high^ monocytes (**Figure 5F**). These results indicate that the HBP pathway contributes to the glucose-dependent regulation of CSF1R expression and monocyte development but is not essential.

## Discussion

The use of blocking Ab-based strategies and genetic models aiming at investigating the role of CSF1R on blood monocyte subsets survival and generation have generally favoured the view that there is a selective impact on the non-classical subset ^20-22,24,26-28^. Non-classical monocytes in rats, the dominant subset in this species, also rely on CSF1R for their differentiation ^44-46^. A humanized anti-CSF1R antibody, axatilimab, has been approved for the treatment of chronic graft versus host disease ^47^. Treatment involves intermittent administration every 2 weeks, which leads to cyclical loss and recovery of CD16^+^ non-classical monocytes. Patients with dominant CSF1R coding mutations have a selective loss of non-classical monocytes^48^. While the effect of CSF1R blockade on mouse Ly6C^low^ monocytes appears consistent and clear, the underlaying molecular and cellular mechanisms leading to the decrease of these cells are not well-defined.

Our analysis indicates that CSF1R signalling is required for the differentiation of blood monocytes. The differentiation of mouse monocyte subsets involves complex interactions amongst several transcription factors. Monocyte development strictly requires Zeb2, which is regulated by C/EBP transcription factors ^49^. In particular, the transcription factor C/EBPα is required for the generation of Ly6C^high^ monocytes but surprisingly the effect of loss of C/EBPα is bypassed to give rise directly to Ly6C^low^ monocytes ^50^. Ly6C^low^ monocyte development involves Notch signalling and depends on the transcription factors RBP-J, Nr4a1 (Nur77), C/EBPβ, BCL6 and IRF2 ^50-53^. Several of these transcription factors bind to multiple regulatory elements in the *Csf1r* locus, whilst exclusively IRF8 binds to FIRE ^54^ and IRF8 regulates monocytopoiesis^55,56^.

A recent study demonstrated that endothelial cell derived CSF1 is required to sustain blood Ly6C^low^ monocytes in a CX3CR1-dependent manner ^23^. Ly6C^low^ monocytes in Tie2^cre^ x CSF1^fl/fl^ mice are almost completely absent from the blood circulation ^23^. We observed decreased Ly6C^low^ monocyte frequencies in CX3CR1^creERT2^ x R26^TdTomato^ x Csf1r^flox/flox^ mice (**Figure 4B**). Neither finding strictly indicates a role for CSF1R in monocyte survival. Depletion or dysregulation of CSF1R-dependent macrophages in any organ could increase extravasation of Ly6C^high^ monocytes reducing their effective half-life in blood. For example, intestinal macrophages, which are replaced continuously by monocytes, are CSF1-dependent^57^.

In view of the impact of CSF1R inhibition, the increased proportion of Ly6C^low^ monocytes in FIRE/FIRE mice compared to littermate controls was unexpected (**Figure 4**). Previous evidence indicated CSF1R was almost undetectable in blood monocytes from these mice. However, we detected residual CSF1R expression on Ly6C^low^, but not Ly6C^high^ blood monocytes in FIRE/FIRE mice. The selective depletion of this subset in FIRE/FIRE mice treated with AFS98 confirmed that despite the low expression the Ly6C^low^ subset remains CSF1/CSF1R dependent. Ly6C^high^ monocyte depletion (by administration of the CCR2-depleting MC-21 Ab) was reported to increase the lifespan of Ly6C^low^ monocytes ^32^. This was paralleled by decreased detection of CSF1R on the remaining Ly6C^low^ monocytes ^32^. We suggest that a similar mechanism underlies regulation in the FIRE/FIRE mice; the absence of the receptor in Ly6C^high^ cells increases the availability of the ligand to the Ly6C^low^ cells. This conclusion leaves open the question of precisely how Csf1r transcription is initiated to enable expression in the absence of FIRE. More generally, the results also call into question the proposed role of CSF1 in the generation and maintenance of monocytes in bone marrow ^58^.

Previous studies demonstrated that Flt3L^-/-^ mice display a more severe immune cell phenotype in comparison to Flt3-deficient animals ^59,60^. In Flt3^-/-^, but not in Flt3L-deficient mice, CSF1R signaling compensated for Flt3 loss, suggesting that in the absence of one RTK another one could functionally compensate and restore phosphorylation of downstream targets ^59^. An additional report using Flt3^-/-^CSF1R^-/-^ mice established a crosstalk between tissue resident macrophages and DCs ^61^. A recent report demonstrated that humans bearing loss-of-function Flt3L-mutations displayed monocytopenia while neutrophil numbers were only moderately decreased^62^. These data suggest that human monocytes display higher Flt3L-dependence in comparison to mouse monocytes ^62^. The underlaying molecular patterns leading to this observation remain to be defined and might explain crucial parts of monocyte biology that differ among species.

CSF1R signaling regulates glucose uptake and metabolism in monocytes (**Figure 5B-C**)^25^. We showed using genetic models and pharmacological inhibitors, that while PFKFB3-dependent glycolysis and the pentose phosphate pathway were not essential for monocyte CSF1R expression and numbers, blocking the hexosamine biosynthetic pathway with tunicamycin strongly reduced CSF1R expression on blood monocytes ^25^. In the current study we generated and validated genetic depletion of GFPT1 and found that CSF1R expression was marginally decreased, leading to diminished Ly6C^low^ monocyte frequency. However, taken together, this and the previous studies suggest that downstream metabolism of glucose by any of the three alternative pathways does not explain the effect of glucose transport inhibition. An alternative possibility is that the generation of glucose-6-phosphate (G6P) *per se* is essential. 2-Deoxyglucose, which competitively inhibits hexokinase, has been shown to induce ER stress and an unfolded protein response in human monocytes ^63^. There is also a close regulatory relationship between G6P, glycolytic enzymes and the activity of the vacuolar ATPase ^64^, which in turn has been implicated in the control of CSF1R trafficking from the Golgi ^65^. These alternative explanations for the essential role of glucose uptake in monocyte differentiation warrant further investigation.

## Acknowledgments

SI is funded by Institut National de la Sante et de la Recherche Medicale (INSERM), Agence Nationale de la Recherche (ANR-22-CE14-0027-02; ANR-21-CE15-0020-02; ANR-23-CE14-0054-02; ANR-23-CE15-0032-01) and by Fondation de France. JWW was supported by NIH R01AI165553. AG was supported by the Lefoulon-Delalande Foundation and the French government through the France 2030 investment plan managed by the National Research Agency (ANR), as part of the Initiative of Excellence Université Côte d’Azur under reference number ANR-15-IDEX-01. NIH supported GJR (R37AI049653) and JH (K00ActfCA264434).

## Author contribution

AG and SI designed the study. AG and SI wrote the manuscript and prepared the figures with input from all co-authors. AG, JM, ZC, CD, EB, JH, BD, AC, SG, MF, GJ, FNZ, EB, FT, RRG, AB, JGN and SI performed experiments. AG, JM, ZC, JH, EB, JGN and SI analyzed data. DD, GJR, PA, AJ, DAH, JWW and MB provided tools and expertise.

## Conflicts of interest

The authors declare no competing interests.

## Material and Methods

### Experimental models

#### Mouse models

Wild-type C57BL/6J (Jax #000664), CX3CR1^gfp^ (B6.Cg-Ptprca Cx3cr1^tm1Litt/LittJ^, Jax #008451), Clec9a^iCre^ (B6J.B6N(Cg)-*Clec9a^tm2.1(icre)Crs^*/J, Jax #025523), Lyz2^Cre^ (B6.129P2-Lyz2tm1(cre)Ifo/J, Jax#004781) and R26^TdTomato^ (B6.Cg-Gt(ROSA)26Sor^tm9(CAG-tdTomato)Hze^/J, Jax #007909) mice used in this study were maintained under C57BL/6J background and originally purchased from Janvier Labs. CX3CR1^CreERT2^ mice were reconstituted from the cryopreserved sperm provided from Steffen Jung (Weizmann Institute of Science, Rehovot, Israel ^32^) to Gwendalyn J. Randolph. Csf1r^ΔFIRE^ mice ^24^ were provided by Dr. David Hume to Dr. Marc Bajénoff. Gfpt1 floxed mice (57BL/6N-Gfpt1<tm1c(EUCOMM)Wtsi>/H) were obtained from Mary Lyon Center (MRC, Harwell UK). Experimental and control animals were co-housed, and littermate controls were used as often as possible. Animals of mixed sex and similar age were used within each cohort. Since we did not observe any sex-specific phenotypes, we decided to group data from male and female mice together in experiments where mice of both sexes were used. Mice were bred and housed in the Mediterranean Center of Molecular Medicine facility (INSERM U1065, Université Côte d’Azur), the PEMED facility (Pasteur campus, Université Côte d’Azur), the Washington University in Saint Louis animal facility, the University of Minnesota Medical School Research Animal Resources facility, or the Centre d’Immunologie de Marseille Luminy facility. An ambient temperature of ∼20-23 °C was maintained, with a 12/12-hour light/dark cycle and food available ad libitum. Animals were euthanized by cervical dislocation. Animal protocols required for experimentation other than organ collection were authorized by the French Ministry of Higher Education and Research upon approval of the local ethical committee (CIEPAL Azur) at Université Côte d’Azur, and by the Institutional Animal Care and Use Committee (IACUC) at Washington University in Saint Louis and University of Minnesota Medical School.

### Genotyping

DNA was extracted from ear biopsies by incubation with 25mM NaOH/0.2 mM EDTA for 60min at 105°C, followed by addition of an equal volume of 40 mM Tris HCL (pH 5.5). DNA was amplified by PCR using DreamTaq Green PCR Master Mix (2X) (Thermo Scientific). PCR products were visualized on a 3% agarose gel.

### *In vivo* treatments

#### CSF1R blockade

Mice were injected intra-peritoneally with 500 μg anti-CD115 blocking antibody (AFS98, BioXCell cat#BE0213) or isotype control (Rat IgG2a, BioXcell cat#BE0089) as indicated in the corresponding figure legends and were sacrificed 16 h after the last injection for analysis of adipose tissue macrophages.

#### Tamoxifen treatment

CX3CR1^creERT2^ TdTomato reporter mice were treated with tamoxifen dissolved in corn oil (20 mg/mL, 100 µL/mouse/administration) by oral gavage as often as indicated in figure 4A.

### *Ex vivo* treatments

Blood was collected in heparinized tubes by submandibular bleeding. Whole blood was incubated at 37°C for 45 minutes with Brefeldin A (5μg/mL), Monensin (2μM), Cytochalasin D ( or Harringtonine (2μg/mL), then processed for flow cytometry analysis. For the CSF1R recycling assay, whole blood was incubated on ice with PBS containing 100ng/mL recombinant CSF1 for 30 minutes. Blood cells were then centrifuged and resupended using 1.5mL PBS at least 5 times to wash away any unbound CSF1. To allow CSF1R recycling, blood cells were resuspended in RPMI and incubated at 37°C for 20 to 100 minutes. Red blood cells were then lysed and white blood cells were analyzed by flow cytometry to quantify surface CSF1R expression.

### Flow cytometry

Tissues were harvested after cervical dislocation and washed in PBS. Splenocytes were prepared by gently crushing the spleen on a 70μm strainer in flow buffer (PBS containing 1% BSA and 2mM EDTA). Bone marrow cells were prepared by flushing femurs and tibias with flow buffer. Blood was drawn from the submandibular vein and collected in heparinized tubes. Leukocyte counts were quantified using a veterinary hematology analyzer (Exigo H400) or directly on the Cytek Aurora cytometer. Red blood cells were lysed from all single cell suspensions using BD Pharm Lyse lysing solution (BdBiosciences cat #555899). Cells were washed with PBS, washed in flow buffer and then stained. For intracellular staining, cells were fixed and permeabilized using Miltenyi Foxp3 staining buffer (cat #130-093-142). All antibodies were used 1/200. A list of all antibodies used is provided in Supplementary Table 1. Flow cytometry data were acquired using a BD FACS Canto II and a Cytek Aurora cytometer with 5 laser configuration. All analyses, including unsupervised t-SNE analysis, were performed using FlowJo software (Tree Star).

### SCENITH analysis

The method was performed as described in^42^. SCENITH reagents kit (inhibitors, puromycin and antibodies) were obtained from www.scenith.com/try-it and used according to the provided protocol for ex-vivo analysis of myeloid cells. Blood was collected in heparinized tubes by submandibular bleeding. Whole blood was incubated at 37°C for 30 minutes with Puromycin (10µg/mL) and either Control, 2-Deoxy-D-Glucose (100mM), Oligomycin (1µM), 2-Deoxy-D-Glucose + Oligomycin or Harringtonine (2µg/mL). Red blood cells were then lyzed using cold BD Pharm Lyse buffer. Cells were first stained with Live/Dead fixable viability dye, washed, and incubated with Fc Block (2.4G2, BioXcell) before surface staining. Cells were fixed and permeabilized using Foxp3 fixation/permeabilization buffer (Miltenyi) and then stained with anti-Puromycin (Alexa Fluor 647). Cells that were not incubated with Puromycin were used as a negative control for anti-Puromycin signal. Cells that received surface staining but no anti-Puromycin staining (full minus one) were used to measure subset-specific autofluorescence in the AF647 channel, and this background signal was subtracted. Glucose dependence, Mitochondrial dependence and Glycolytic capacity were calculated as previously described ^42^.

### Western Blot

Proteins were extracted from BMDM using a Qiagen AllPrep DNA/RNA/protein Mini kit and subsequently resolved by SDS-PAGE and transferred to PVDF membranes by wet transfer during 90 minutes at 100 V at 4°C. Membranes were blocked in 5% (m/v) powdered non-fat milk in TBS and incubated with primary antibodies (1:1000 rabbit anti-GFPT1 monoclonal EPR4845 (Ab15069 Abcam) and 1:5000 mouse anti-βactin in blocking solution overnight at 4°C. After washing, membranes were incubated with HRP-conjugated secondary antibodies for 1 h at room temperature. Proteins were visualized using a Fusion FX (Vilber Lourmat, France).

### Statistical Analysis

All data are represented in mean ± SEM. Statistical analysis was performed with GraphPad Prism 10 as indicated in each figure legend. Presence of statistically significant differences between conditions are indicated as follows: ns p>0.05; * p<0.05; ** p<0.01; *** p<0.001; **** p<0.0001.

---

**Supplementary Figure 1.**
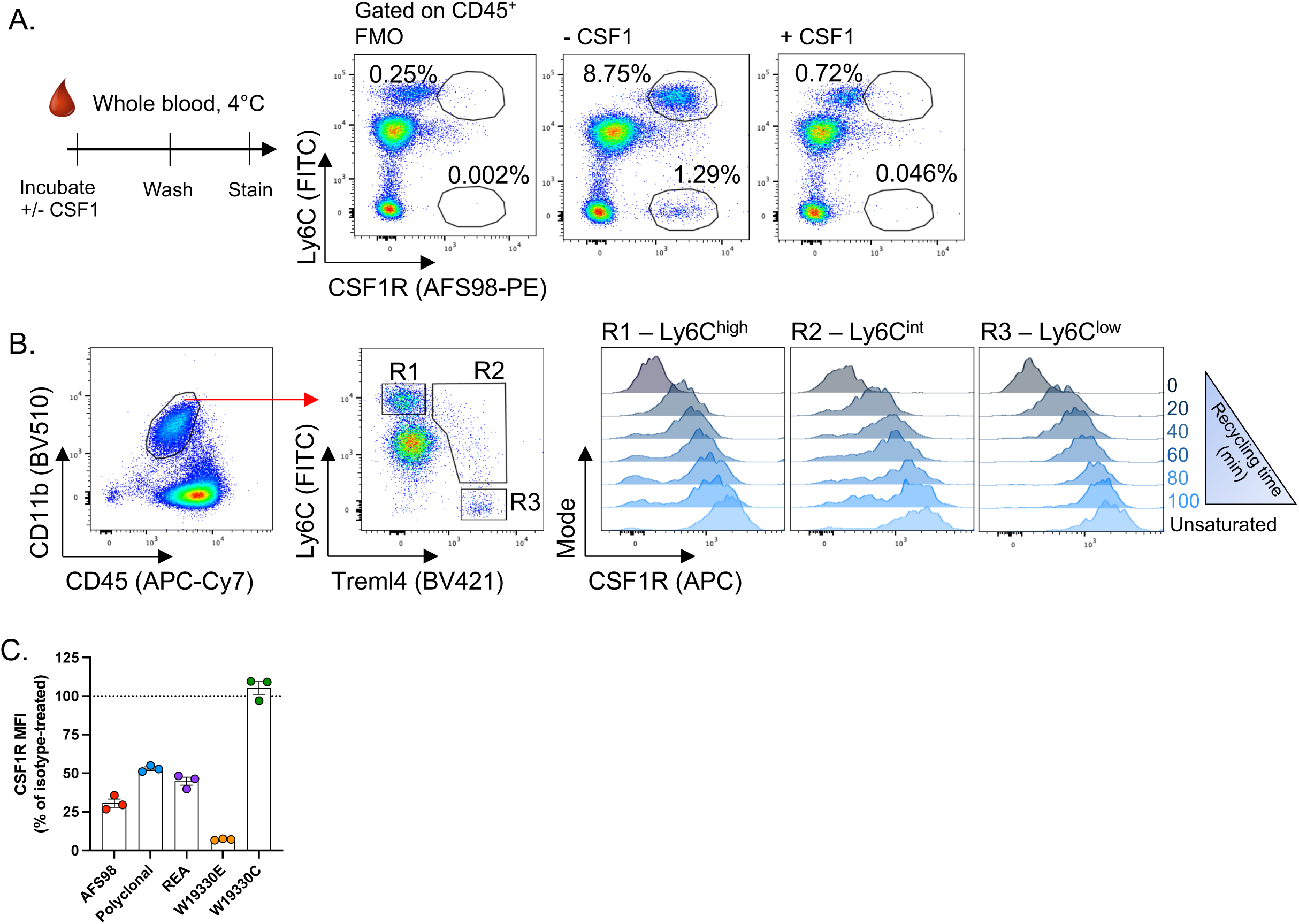
(**A**) Experimental scheme used to test whether epitope masking by CSF1 on CSF1R could occur (left) and flow cytometry plots showing expression of CSF1R on Ly6C^high^ and Ly6C^low^ blood cells depending on CSF1 presence, detected by PE-conjugated AFS98. (**B**) Flow cytometry plots showing CSF1R recycling in blood monocyte subsets. (**C**) Expression of CSF1R measured 24 hours after AFS98 injection using several clones of PE-conjugated antibodies.

**Supplementary Figure 2.**
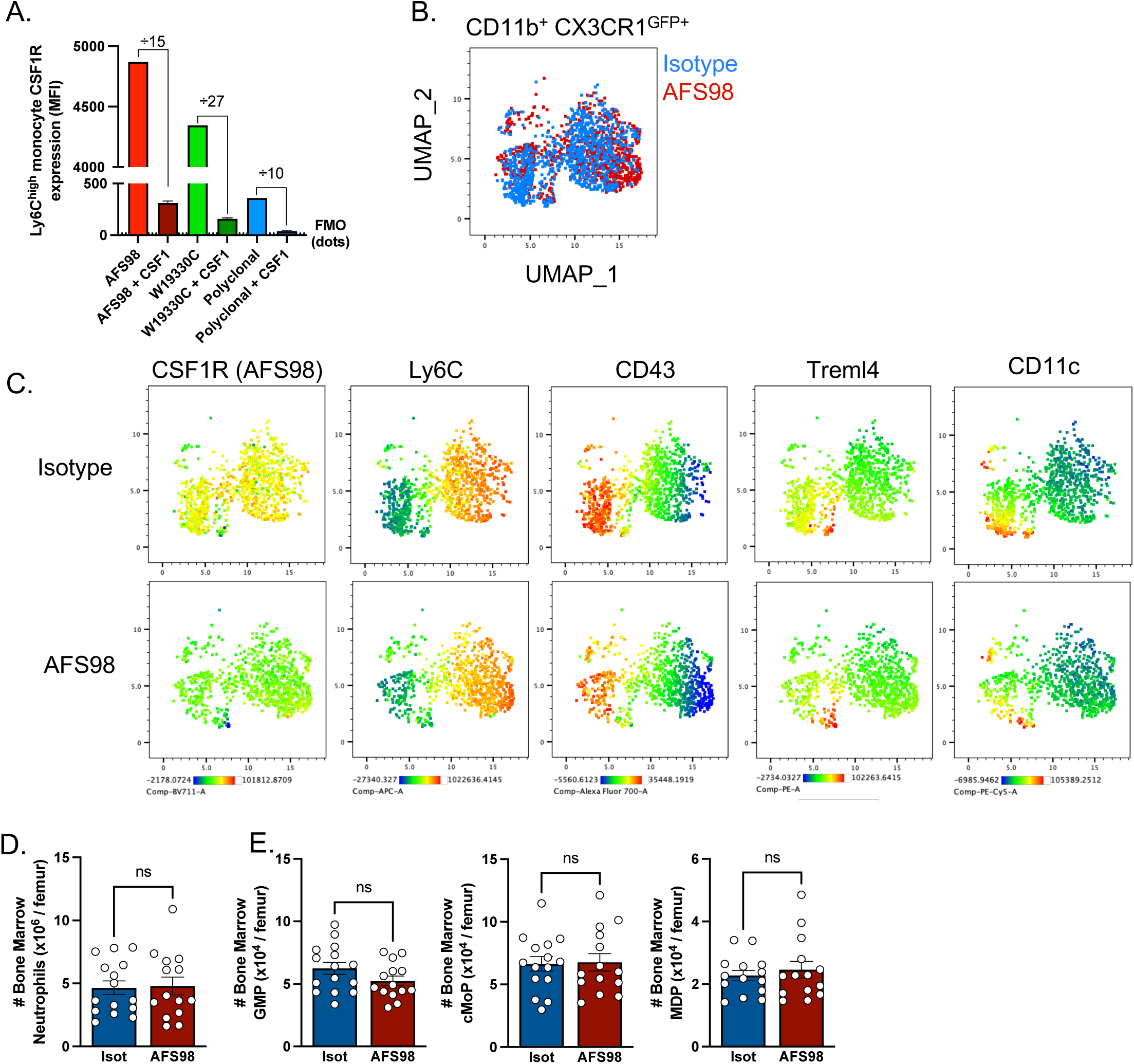
(**A**) Measure of CSF1R expression using AFS98, sheep polyclonal antibodies or clone W19330C on blood monocytes incubated or not with recombinant CSF1. (**B**) Uniform Manifold Approximation and Projection (UMAP) representation of CD11b^+^ CX3CR1^GFP+^ cells concatenated from isotype-treated and AFS98-treated mice. (**C**) UMAP representation of CSF1R (AFS98-BV711 conjugated antibody), Ly6C, CD43, Treml4 and CD11c expression by blood CD11b^+^ CX3CR1^GFP+^ cells from isotype-treated and AFS98-treated mice. (**D-E**) Quantification of bone marrow neutrophils (D) and monocyte progenitors (E) following AFS98 treatment as illustrated in Figure 1A.

**Supplementary Figure 3.**
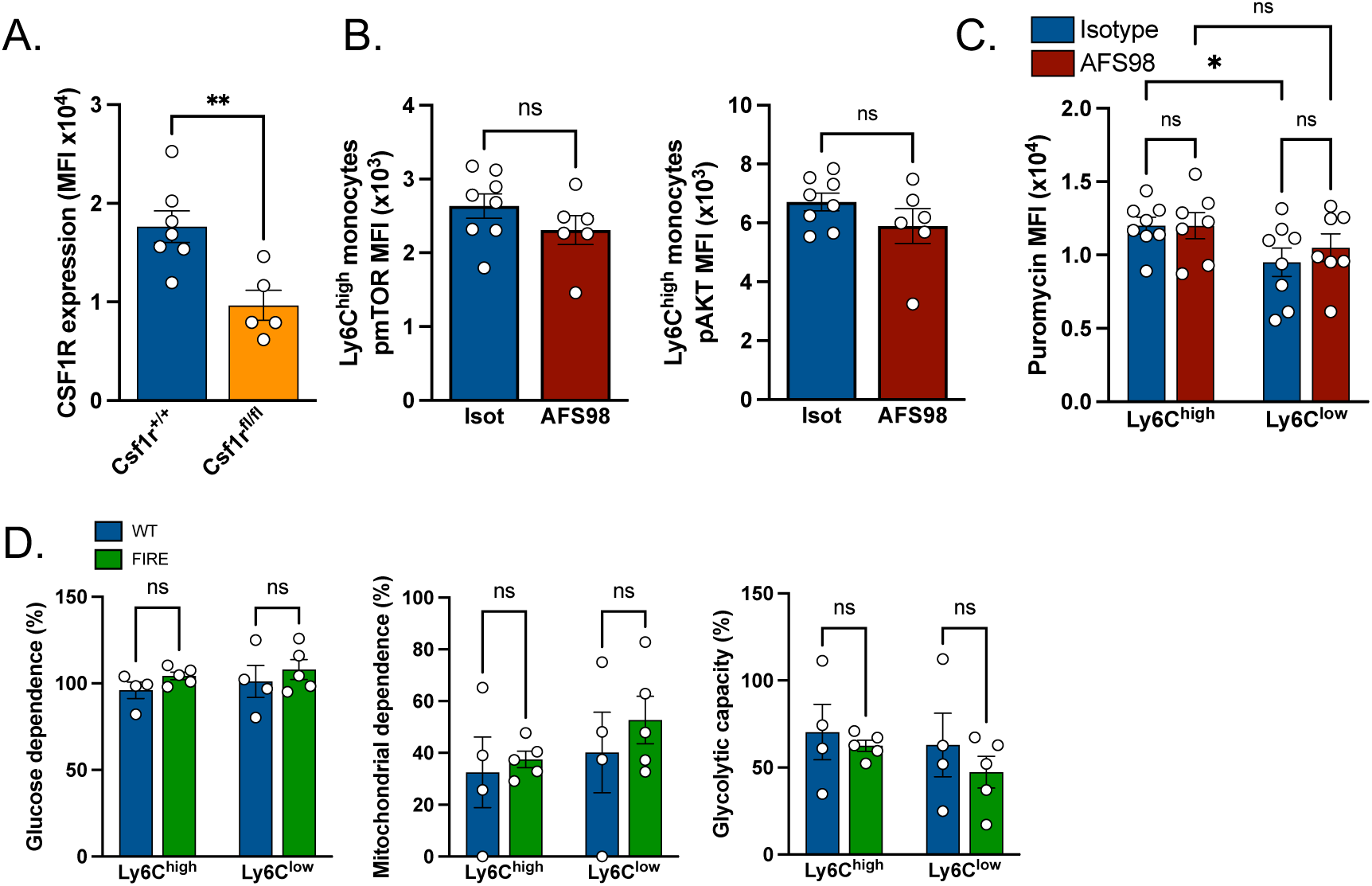
(**A**) Expression of CSF1R on blood monocytes from CX3CR1^creER^ x Csf1r^fl/fl^ mice or Csf1r^+/+^ controls. (**B**) Quantification of mTOR and AKT phosphorylation in blood Ly6C^high^ monocytes following CSF1R blockade or isotype control administration. (**C**) Puromycin incorporation in Ly6C^high^ and Ly6C^low^ blood monocytes in control and AFS98-treated mice. (**D**) Metabolic profile of blood monocytes from Csf1r^ΔFIRE/ΔFIRE^ (FIRE) mice or their littermate Csf1r^ΔFIRE/+^ controls (WT) measured using SCENITH.

**Supplementary Figure 4.**
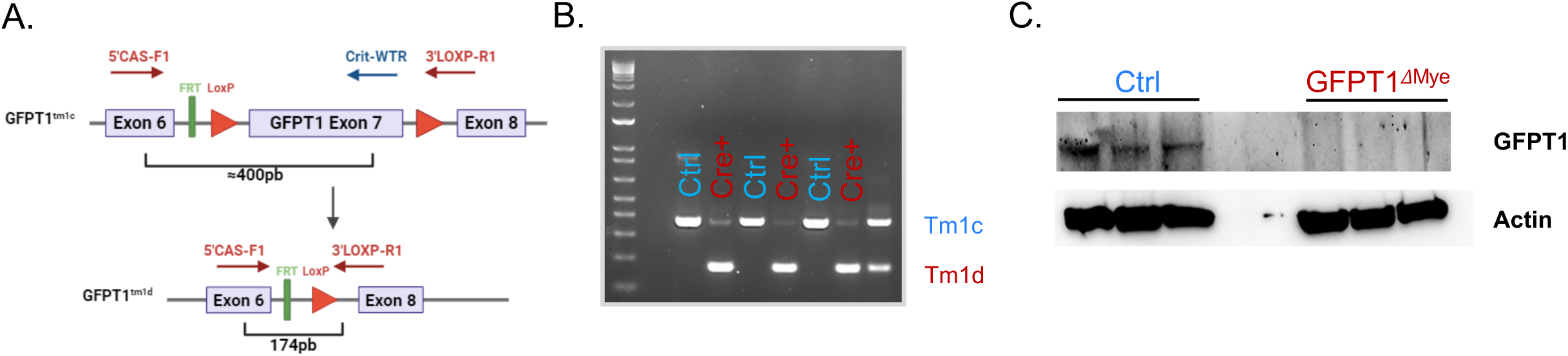
(**A**) Construction of myeloid cell-specific GFPT1-deficient mice. Mice carrying GFPT1 Tm1c allele with exon 7 flanked by LoxP sites (GFPT1 fl/fl) are crossed with Lyz2 Cre mice, resulting in the GFPT1 Tm1d allele in which exon 7 has been excised. **(B)** PCR data on genomic DNA from bone marrow derived macrophages (BMDM) from control (GFPT1 fl/fl) mice or GFPT1^ΔMye^ mice (Lyz2 Cre x GFPT1 fl/fl) mice, showing the presence of Tm1c and Tm1d alleles. As a control, PCR data was shown for an ear biopsy from a GFPT1^ΔMye^ mouse. **(C)** Western blots analysis of GFPT1 expression. Protein extracts were obtained from BMDMs from control and GFPT1^ΔMye^ mice. Beta Actin was used as a loading control.

**Supplementary Table 1.**
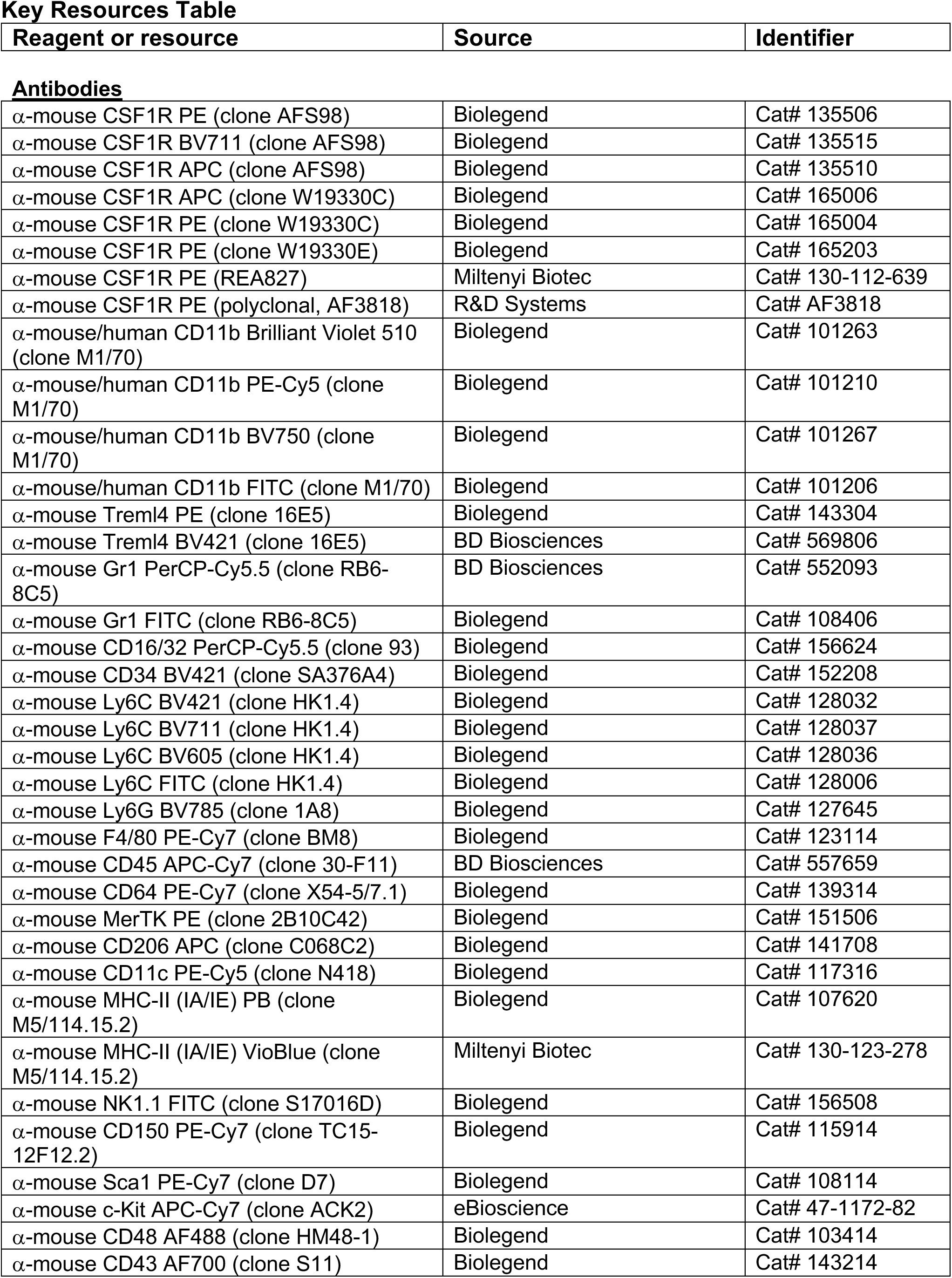
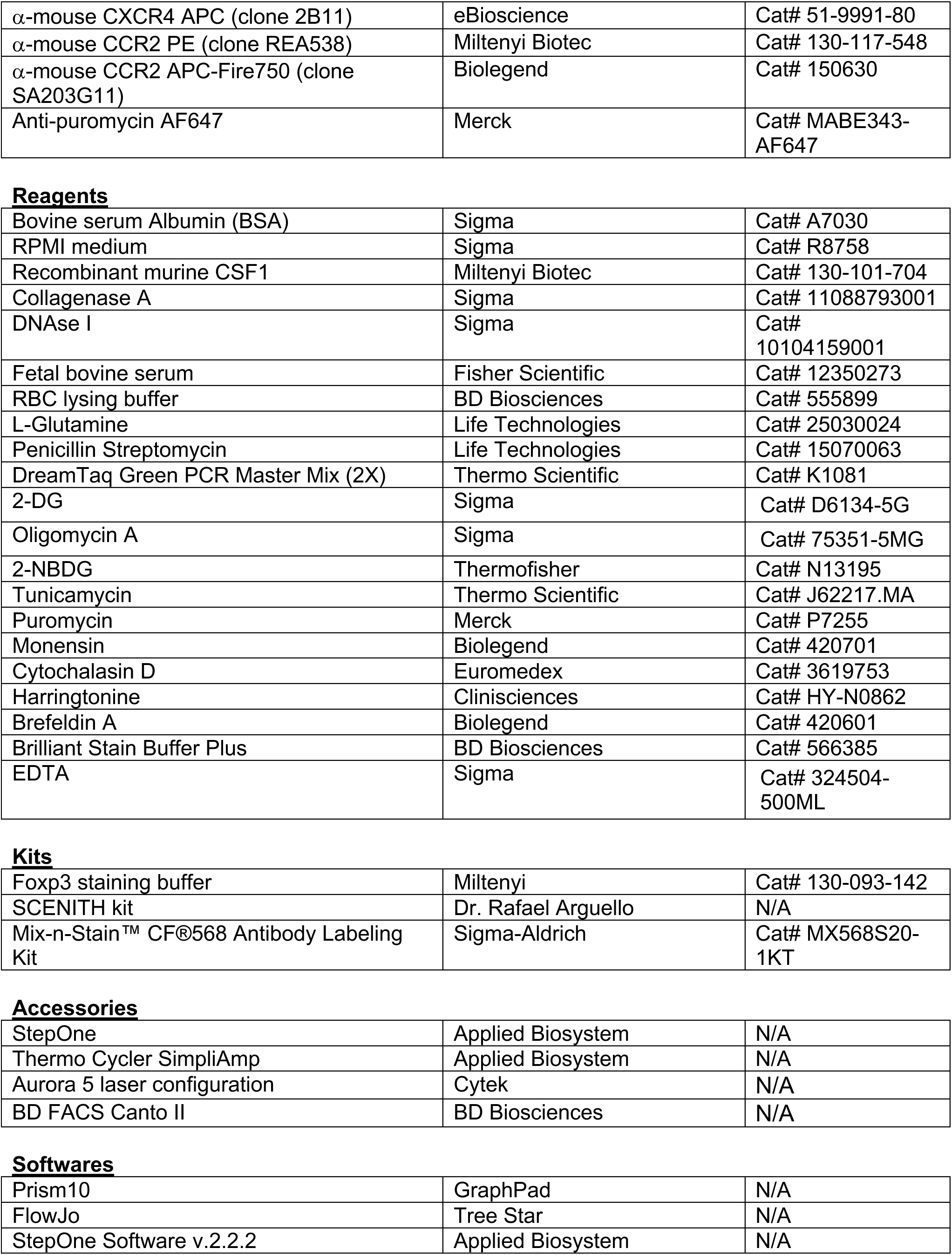

## Notes

### Competing Interest Statement

The authors have declared no competing interest.

